# Deep Learning Enable Untargeted Metabolite Extraction from High Throughput Coverage Data-Independent Acquisition

**DOI:** 10.1101/2020.03.22.002683

**Authors:** Hongchao Ji, Hongmei Lu, Zhimin Zhang

## Abstract

The sequential window acquisition of all theoretical spectra (SWATH) technique is a specific variant of data-independent acquisition (DIA), which is supposed to increase the metabolite coverage and the reproducibility compared to data-dependent acquisition (DDA). However, SWATH technique lost the direct link between the precursor ion and the fragments. Here, we propose a deep-learning-based approach (DeepSWATH) to reconstruct the association between the MS/MS spectra and their precursors. Comparing with MS-DIAL, the proposed method can extract more accurate spectra with less noise to improve the identification accuracy of metabolites. Besides, DeepSWATH can also handle severe coelution conditions.

---

Data dependent acquisition (DDA) selects single precursor ion for fragmentation each time, which has the direct link between the precursor ion and its fragments. In contrast, data independent acquisition (DIA) often uses a wide isolation window (10 Da – 25 Da) for precursor ions selection. It allows a full coverage of observable molecules but at the expense of losing the direct link between the precursor ion and the fragments. Therefore, how to establish the link is a fundamental problem when processing DIA dataset.

In proteomics, methods for this problem can be divided into two categories: peptide-centric methods and spectrum-centric methods. Peptide-centric methods usually need experimental or in silico spectral database^1–3^ of all known peptides for a specific biosystem. Basing on the known spectra, OpenSWATH^4^ uses reverse spectrum matching to locate the targeted peptides and precursor-fragment elution curve correlation to score the confidence of the extracted MS/MS. Specter^5^ can also use curve resolution to deconvolve the multiplexed MS/MS spectra. Some peptide-centric methods, such as PECAN^6^ and DIA-NN^7^, can apply the peptide sequence directly to the MS/MS deconvolution. Spectrum-centric methods, such as DIA-Umpire^8^ and Group-DIA^9^, detect covarying precursor-fragment group and generate pseudospectra from DIA. Therefore, spectrum-centric methods are spectral library-free methods.

Analogously, data analysis methods for DIA-based metabolomics can be also categorized into metabolite-centric and spectrum-centric methods. Since it is difficult to infer accurate m/z values of fragments from the structures of metabolites, metabolite-centric methods, such as MetDIA^10^ and MetaboDIA^11^, rely on DDA-based metabolite spectra library. However, the spectral library is limited by the available of metabolite standards. Because of this, metabolite-centric methods can’t take fully advantage of the coverage power of DIA. Spectrum-centric methods can resolve MS/MS spectra of metabolites from DIA data, which is preferred for untargeted metabolomics. DecoMetDIA^12^ applied the hierarchical clustering strategy to reconstruct the pseudo-spectra of metabolites from the co-eluted of fragments. MS-DIAL^13^ resolve the MS/MS spectrum from multiplexed spectra by least squares. Both DecoMetDIA and MS-DIAL try to solve the MS/MS reconstructing problem in DIA-based metabolomics. However, the hierarchical clustering of DecoMetDIA depends heavily on the quality of peak shapes. Co-eluted and irregular peaks will deteriorate the clustering result. Background ion and random noise will weaken the effectiveness of the least-squares based deconvolution in MS-DIAL and introduce the impure ions into the reconstructed MS/MS spectra.

Here, we propose DeepSWATH, a spectrum-centric method for untargeted metabolite extraction from SWATH-MS dataset based on deep neural network. The core of DeepSWATH method is the precursor-fragment correlation (PFC) model to extract the fragments of a precursor ion. It is a fully untargeted extraction method. Once the model is trained, spectra library will not be need when processing a new dataset.

PFC model was trained by a combination dataset including both experimental data and simulated data. Experimental data were obtained from a large open access metabolomics dataset, and its MetaboLights identifier is MTBLS417. Twenty healthy control samples of DDA dataset were processed by XCMS to obtain the golden standard of the precursor-fragment groups. The extracted ion chromatograms (EIC) of the precursors and fragments were generated from the corresponding DIA dataset, regarded as positive samples. At the retention time of the precursor ion chromatographic peak, the other fragment ions also exist besides the fragment ions. We randomly chose the equal number of these fragment ions (decoys), extracted the EICs and took them as negative samples (Figure 1). In order to handle coelution condition well, we also generate a number of simulated samples, because coelution is relatively infrequent in the experimental data. Each simulated sample includes a Gaussian like precursor profile and a correlated or uncorrelated fragment/decoy profile.

**Figure 1.**
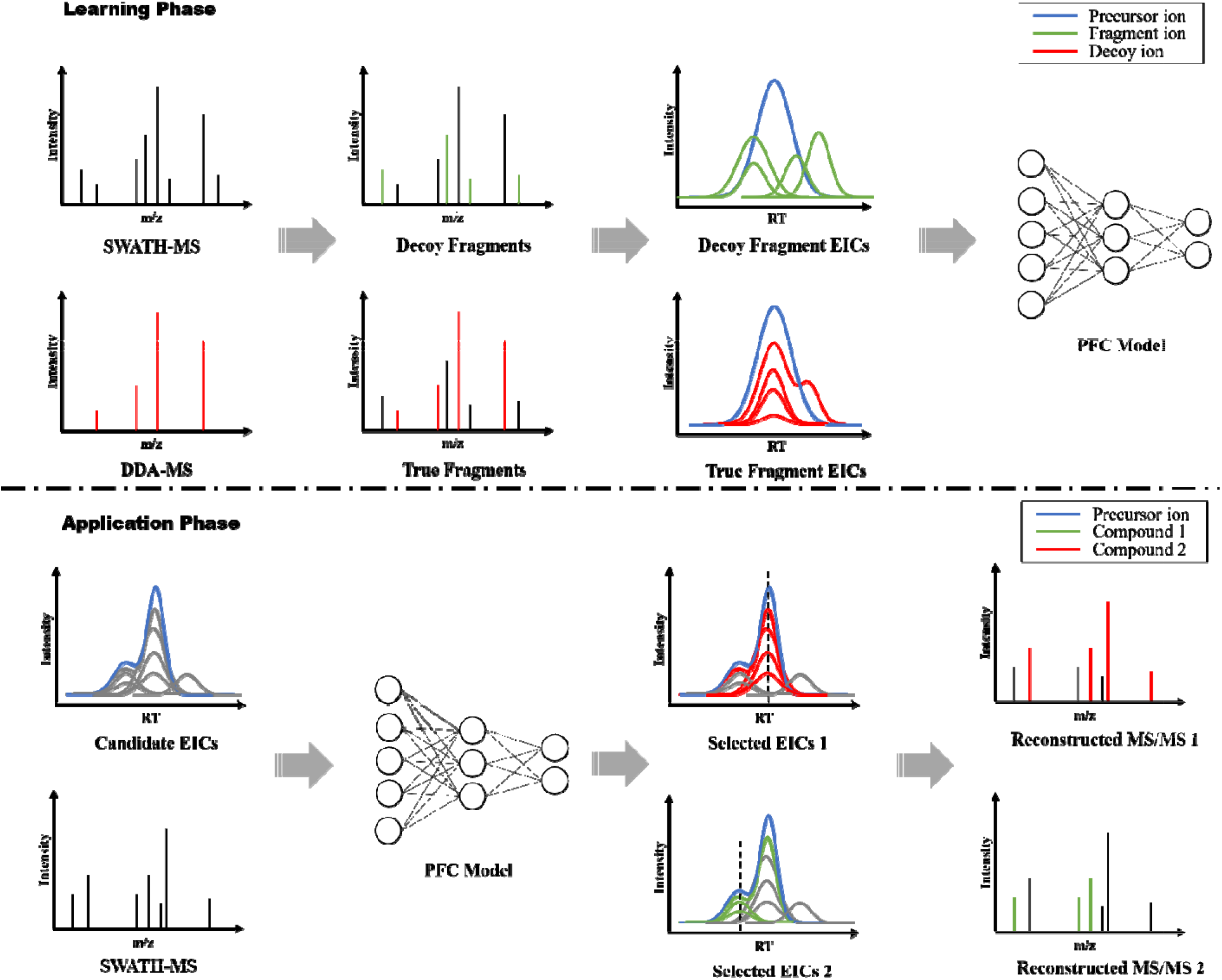
Workflow of DeepSWATH. Training data were obtained by a DDA-DIA compared dataset. *Training P ase*: Ground truth of MS/MS were extracted from DDA data. Targeted extraction of fragmental ions was performed on the corresponding DIA data. The elution curves of fragmental ions together with the elution curves of their precursors were treated as positive data. Decoy ion elution curves were random chosen from noise, background or interferential ions. They were treated as negative data. The data were used for training PFC Model. *Application Phase*: With the PFC model, when given another DIA data, the MS/MS can be reconstructed by: 1. Extracting precursor features with XCMS, MZMine or any other software; 2. Extracting all candidate fragmental EICs of each precursor; 3. Predict the true fragmental ions of each precursor with the PFC model.

In the learning phase, the input is the precursor EICs and the corresponding fragment/decoy EICs, followed by a series of hidden layers. The output layer is a binary classification layer (Figure 2). In the application phase, precursor EICs of the new DIA dataset can be extracted by any LC-MS data processing framework, including XCMS^14^, MZMine^15^, OpenMS^16^ or KPIC2^17^. DeepSWATH will extract the EICs of all possible precursor-fragment pairs and predict the relationship between precursor-fragment pairs with the trained PFC model. The MS/MS spectra can be reconstructed with the prediction results easily. Detailed methods are described in *methods* section.

**Figure 2:**
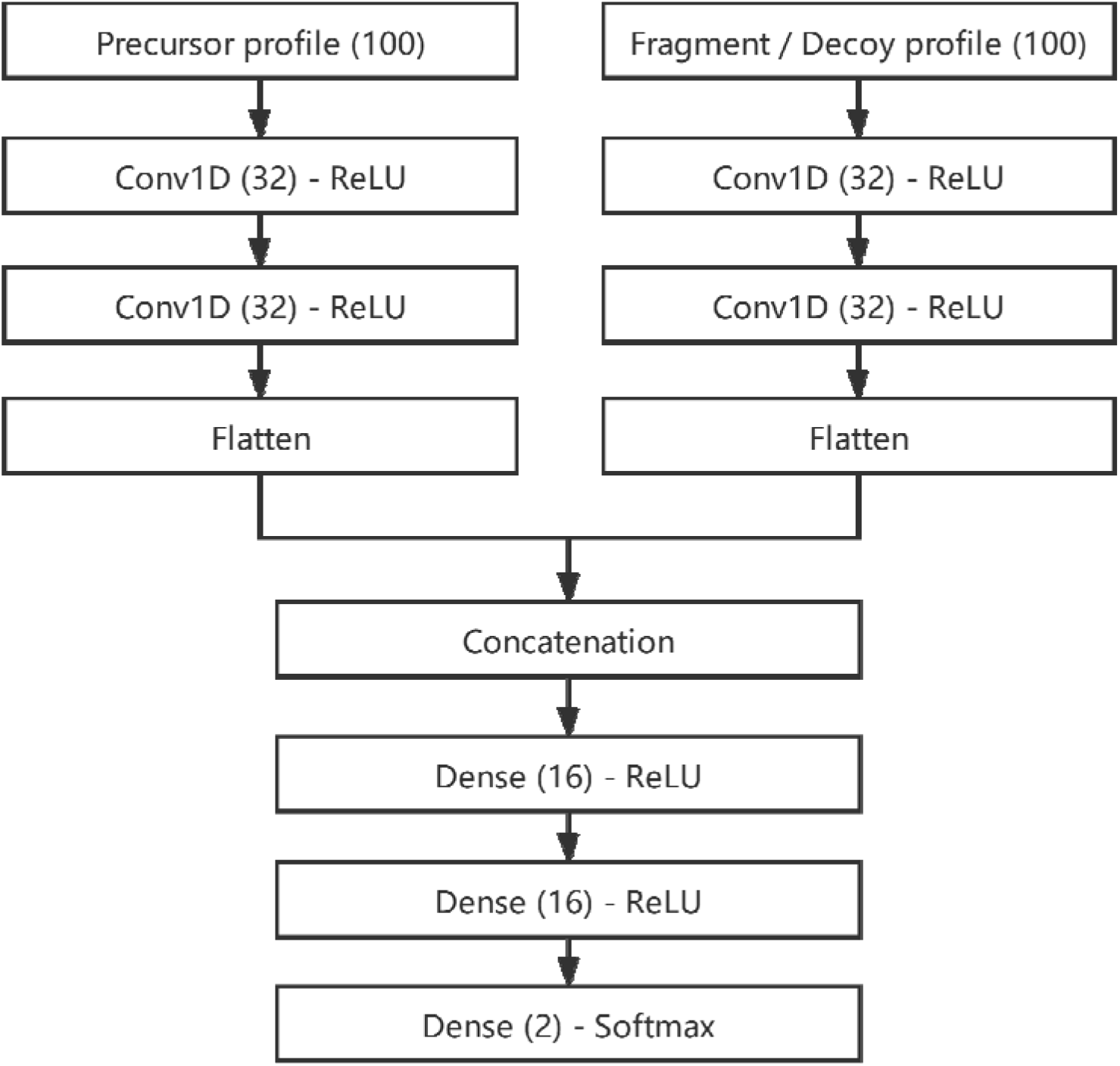
The structure of the neural network. The input precursor profile and fragment/decoy profile were followed by two convolution layers, respectively. Then, the outputs of the convolutional layers were flattened, concatenated and followed by two dense layers. The output layer is a binary classifier, which predicts the input pair of precursor-fragment is associated or not.

## Results

### Deep learning-based PFC model

The PFC model is a precursor-fragment relationship classifier which can be regarded as a binary classifier to differentiate true fragment ion of a precursor from the interferential ions. The deep neural network of PFC model was implemented using Keras with Tensorflow backend. The precursor and fragment/decoy EICs from MTBLS417 were randomly split into three subsets: an 80% subset for training (training set), a 10% subset for optimizing the hyper parameters (validation set), and a 10% subset for evaluating the performance (test set).

During the learning phase, the classification accuracy and loss was used to monitor the performance of the model. The training procedure was stopped after 15 epochs because of no obvious improvement of the loss and accuracy. A callback function was used to reduce the learning rate dynamically by monitoring the loss of the validation set. The test set was used to evaluate the performance of the PFC model, the precision, recall and accuracy were 0.930, 0.936 and 0.933 respectively. The area under the ROC curve is 0.98. The results indicate that the PFC model can effectively differentiate whether the EICs of the given precursor and fragment are associated or not.

### Accuracy of Reconstructed MS/MS

We evaluated the accuracy of reconstructed MS/MS with two datasets. One is a mixture sample with 30 standard compounds (30 STD)^10^, the accuracy was evaluated by comparing the reconstructed MS/MS spectra with the spectra in database. The other one is the case group of MTBLS417, the accuracy was evaluated by comparing the reconstructed MS/MS spectra with the spectra acquired by DDA. The criterion is the Pearson correlation coefficient between the reconstructed MS/MS spectra and the reference spectra.

Figure 3(A) and 3(B) are the distribution of Pearson correlation coefficients of DeepSWATH and MS-DIAL on 30 STD and MTBLS417 dataset. For the 30 STD dataset, the median values of DeepSWATH and MS-DIAL are 0.9928 and 0.9916, respectively. For the positive mode of MTBLS417 dataset, the median values of DeepSWATH and MS-DIAL are 0.8723 and 0.8669, respectively. For the negative mode, the median values of DeepSWATH and MS-DIAL are 0.8524 and 0.8353, respectively. The detailed results generated by MS-DIAL and DeepSWATH can be found in Table S1 – S10.

**Figure 3.**
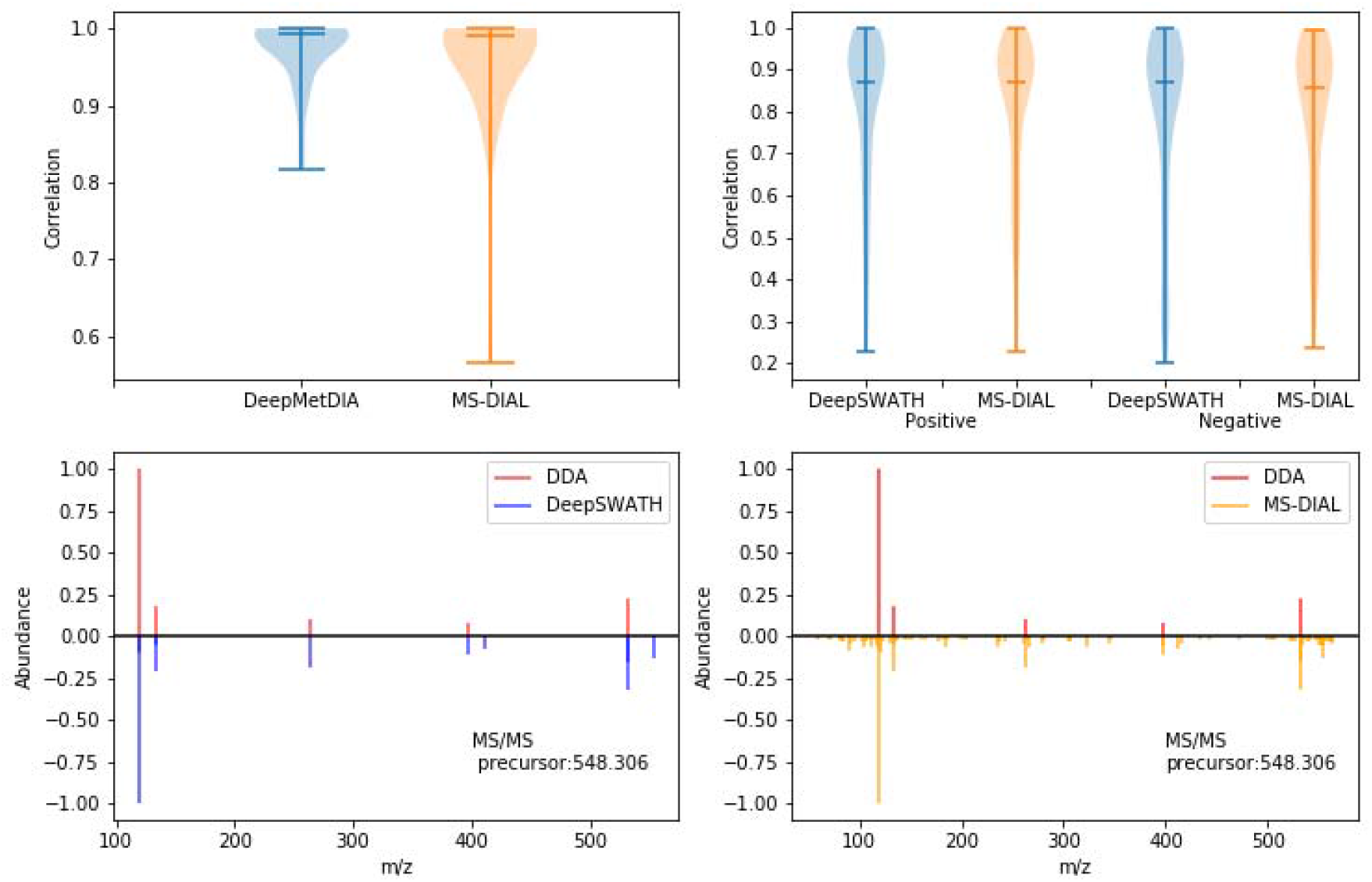
The distribution of the MS/MS reconstruction accuracy for 30 STD dataset (A) and MTBLS417 dataset (B), with the y-axis is the Pearson correlation between the reference MS/MS and the reconstructed MS/MS. Comparison with an example between DDA spectrum and reconstructed MS/MS obtained by DeepSWATH (C) and MS-DIAL (D).

Figure 3(C) and 3(D) is an example of the reconstructed MS/MS spectra. Both DeepSWATH and MS-DIAL can resolve the main fragment ions. However, DeepSWATH is more robust to interferential ions and noise when compared with MS-DIAL, and the reconstructed MS/MS is cleaner. It is important to obtain clean MS/MS spectrum for metabolite identification, because the presence or absence of fragments has large influence to the value of scoring function in many in-silicon methods. For example, SIRIUS^18,19^ takes fragmentation trees as important features to predict substructures of metabolites, which also takes all the fragments into consideration.

### Significance of cleaner MS/MS spectrum

In order to justify the cleaner MS/MS spectrum obtained by DeepSWATH can authentically improve the metabolite identification, As shown in Figure 4, MS/MS spectra of DDA and reconstructed spectra by DeepSWATH and MS-DIAL were imported into SIRIUS (version 4.0.1) for identification. In the first example, the Pearson correlations of DeepSWATH and MS-DIAL are 0.822 and 0.807, respectively, while in the second example the correlations are 0.769 and 0.838, respectively. From the results given by SIRIUS, the identified structures of DDA and DeepSWATH spectra are consistent. However, with the interferential peaks in the MS-DIAL spectra, SIRIUS gave different results from DDA spectra, although the correlation of the second example is even higher than DeepSWATH. The results indicate that the cleaner MS/MS spectra obtained by DeepSWATH is of significance in the metabolite identification.

**Figure 4.**
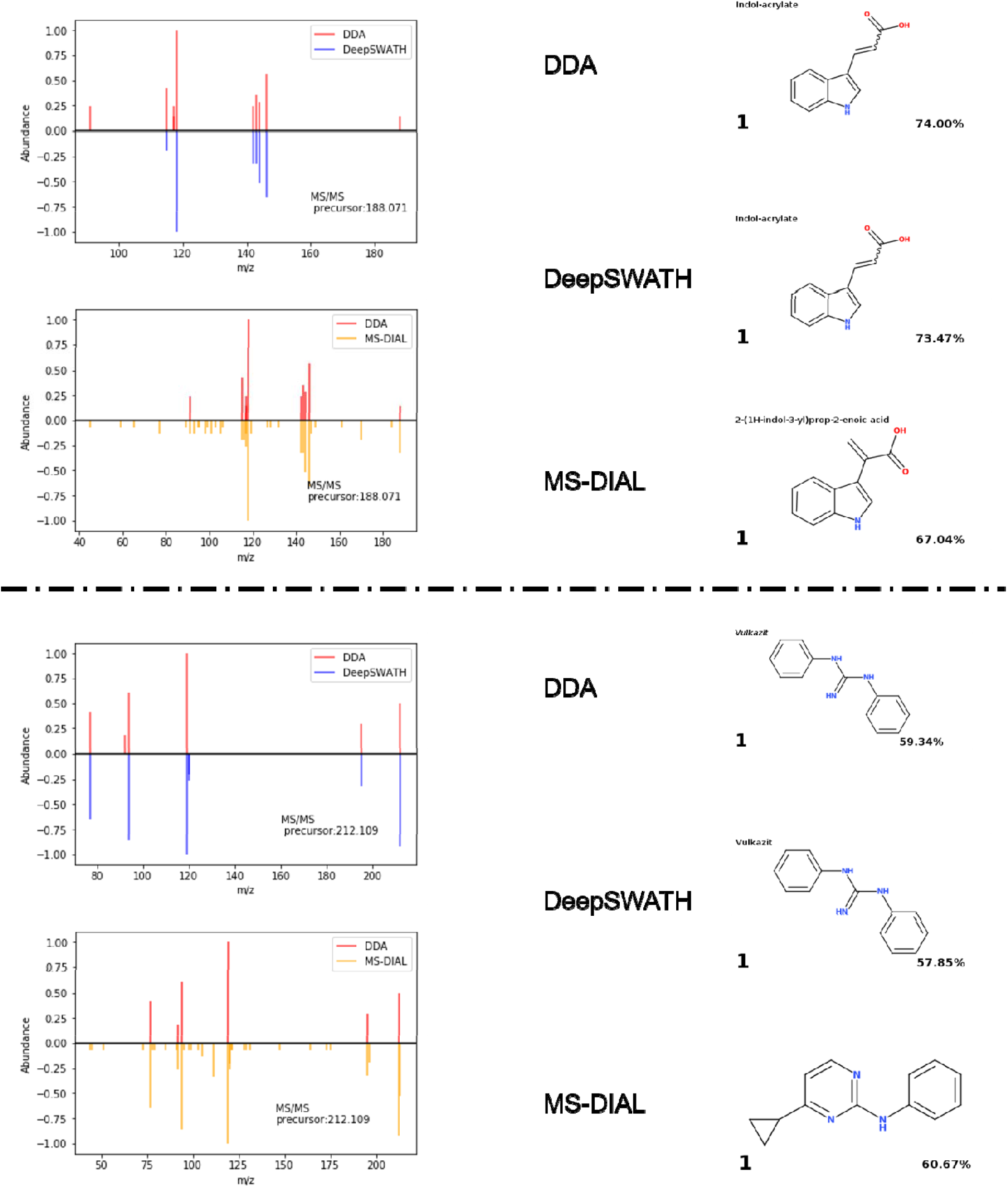
Comparison of DDA spectra and reconstructed spectra by DeepSWATH and MS-DIAL (left), and the corresponding identification results given by SIRIUS (right). The percentage number at the lower right of the annotated structure indicate the confidence coefficients.

### Coelution handling

To evaluate the ability of DeepSWATH in handling the coelution problem, two examples were shown in Figure 5. In the first example, the two precursors have the same m/z value, which is 266.094. Their retention times are also similar, and their chromatographic peaks are overlap (A1). After processed by DeepSWATH, the fragment ions were successfully assigned to the appropriate precursor ions, which indicates that the coelution of precursors did not affect the accuracy of MS/MS reconstruction (A2 and A3). In the second example, the EIC of the precursor includes a single peak, while several fragment ions show are coeluted (B1). It means the precursors share some fragments with the same m/z values. From the results, it can be seen that the true fragments were also successfully assigned (B2). If a higher accuracy of the fragment abundances is needed, it can be combined with multivariate curve resolution (MCR)^20^ to resolve the common fragments. With MCR-ALS, the pure concentration of each fragments can be obtained (B3). Therefore, DeepSWATH is also effective under the coelution situation.

**Figure 5.**
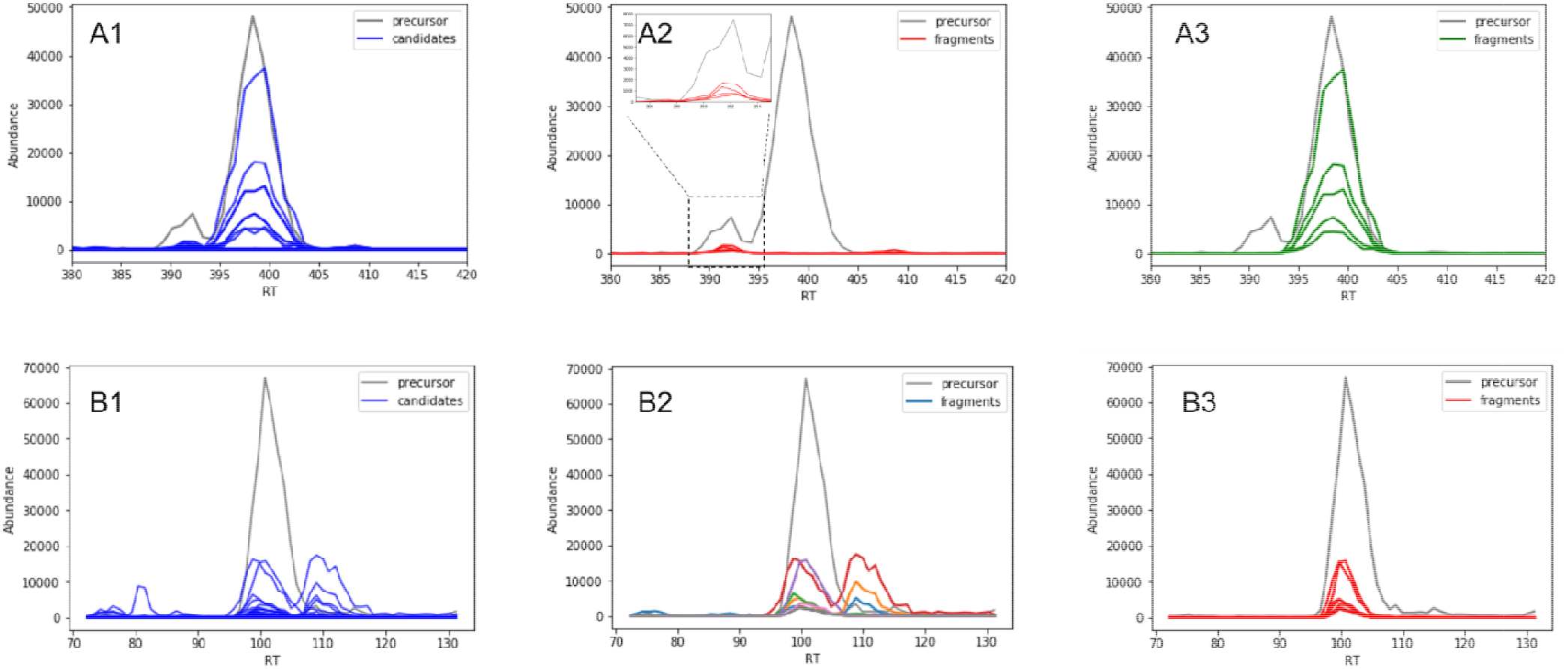
Examples of coelution handling of DeepSWATH. In the first row, the EICs show coeluted precursors and standalone fragments (A1). DeepSWATH assigned the corresponding fragments to the precursors successfully (A2, A3). In the second row, the EICs show a single precursor and standalone fragments, coeluted fragments and interferential fragments. DeepSWATH can assign the true fragments, both standalone and coeluted to the precursor (B2). If needed, it can also combine with MCR-ALS for resolving the coeluted fragments for a more accurate abundances (B3).

## Discussion

In summary, DeepSWATH provides a deep-learning approach to reconstruct MS/MS spectra from SWATH-MS-based metabolomic data. A precursor-fragment model was trained with publicly available dataset, which can predict the relationship between the precursor ion and the fragment ions. Given a new dataset, MS/MS of each precursor can be reconstructed easily with the assistance from this model. We evaluated the performance of DeepSWATH with mix standard dataset and human serum dataset. The reconstructed MS/MS were compared with the spectra in standard database and spectra obtained from DDA. We found that the MS/MS obtained by DeepSWATH is more accurate than the MS/MS obtained by MS-DIAL and can achieve the consistent metabolite identification results with DDA MS/MS spectra. In addition, DeepSWATH can also handle coelution condition.

## Methods

### Experimental training data

The experimental training data were constructed from the healthy control group of MTBLS417 dataset, which analyzed human serum in both DDA and DIA mode. First, we processed the dataset with XCMS package (version 3.8.1). The DDA and DIA data files were processed with the CentWave method and the parameters were the same: snthresh = 10, noise = 200, ppm = 30 and peakwidth = (5, 60). Second, we compare the features obtained from DDA and DIA data. If a feature were detected both in DDA and DIA data (m/z difference < 0.05 Da and RT difference < 30 s), we matched the fragments of DDA with DIA. The fragments obtained from both DDA and DIA were treated as true fragments, and the fragments obtained only from DIA were treated as decoy fragments. Third, we extracted the ion traces (EICs) of the true fragments and their precursors from the DIA data. Besides, we also random selected the same number of decoy ions and extracted their traces. The m/z bin size of the EICs were 0.1 Da and the length of the EICs were 30 s. Finally, we used a linear interpolation to unify the retention times of precursor, fragment and decoy EICs.

### Simulated training data

We added the same number of simulated samples into the training set as the experimental samples to handle the coelution condition. The reason is that the coelution is supposed to be relatively unusual, but has to be under consideration. In the simulated samples, the precursor-fragment pairs involve four conditions: 1. Standalone precursor ion and fragment ion; 2. Coeluted precursor ion and standalone fragment ion; 3. Standalone precursor ion and coeluted fragment ion; 4. Coeluted precursor ion and fragment ion. They were appeared in the simulated dataset randomly. Standalone ions mean each profile includes a single peak and coeluted ions mean each profile includes overlapped peaks. All of the peaks are Gaussian peaks with random sigma values between 1 and 10. The true peaks are in the middle of the EIC vectors, while the coeluted peaks are with a random distance to the true peak. The fragment ions have the same peak position as the precursor ion, while the decoy ions have a different but random peak position.

### Network structure of PFC model

We trained the PFC model based on a deep neural network classifier. Since the EIC is a continuous curve, we chose convolutional neural network as basis. The EICs of precursor ion and fragment/decoy ion are used as input of the PFC model to predict the relationship between them. Therefore, we used a network similar to the text similarity model, which is commonly adapted to natural language processing for synonym analysis. The neural network structure is two convolutional layers followed by the input EICs of precursor and fragment/decoy ions, respectively, and two fully-connected layers followed by the concatenating layer. The output layer is a SoftMax function for classification. All hidden layers employ the ReLU function as activation function. Parameters in the neural network were optimized by the Adam optimizer with initial learn rate as 0.001, and the loss function was set as categorical cross-entropy.

### MS/MS reconstruction with DeepSWATH

The evaluation datasets in this work were the mix standard compounds dataset (30 STD) and the case group of the MTBLS417 dataset. First, the datasets were both processed by XCMS to extract the features of the precursors. For a wider coverage, we employ a looser parameter sets: snthresh = 5, ppm = 100 and peakwidth = (5, 60) to extract the EICs of the precursor ions and all of the possible fragment ions (candidate ions). The m/z bin size of the EICs were 0.1 Da and the length of the EICs were 30 s. Similarly, the linear interpolation was also used to unify retention time of the EICs of the precursor ion and the candidate ions. Finally, the precursor ion EICs and the candidate ion EICs were used as the input. The PFC model was used to predict whether a candidate ion is a fragment of the precursor or not. With the predicted results, the MS/MS spectra were reconstructed easily.

### Combination with MCR-ALS

Sometimes, one coeluted fragment can be assigned to multiple precursors, which means there are multiple precursors share the same fragments. Although it does not affect the results of fragment presence or absence, it may affect the relative abundance of fragments. Thus, DeepSWATH and MCR-ALS were combined together to resolve the coeluted fragments for more accurate abundances. First, the EICs of all predicted fragments were extracted and combine into a m*n matrix, where *m* is the number of the scans of an EIC and *n* is the number of fragments. Second, evolving factor analysis (EFA) was used for estimating the initial concentration curves. Third, alternate least squares (ALS) algorithm was used for resolving the pure chromatograms of all components, in which the main component is the supposed precursor. Finally, fragments with better abundance can be obtained.

### MS/MS reconstruction with MS-DIAL

MS-DIAL software (version 4.12) was used to process the same datasets for comparison. The files in mzXML format were convert to ABF format with the ABF file converter (version 4.0.0) for MS-DIAL. The parameters were set as: minimum peak height = 500, mass slice width = 0.1 and sigma window value = 0.5. The other parameters were kept default. Finally, the results were export in MSP format. The MSP files were then parsed with the functions in libmetgem package^21^.

### Evaluation criterion

The accuracy of MS-DIAL and DeepSWATH were evaluated by comparing the reconstructed MS/MS spectra with the reference MS/MS spectra. For the 30 STD dataset, the reference MS/MS spectra were from the standard database according to the MetDIA paper (only top 5 peaks were given). For the MTBLS417 dataset, the reference MS/MS spectra were obtained from the corresponding DDA files which were extracted by XCMS package. The reconstructed MS/MS spectra and the reference MS/MS spectra were converted to vectors by binning the intensity values in 0.1 Da intervals. Finally, the correlation coefficients were calculated using the Pearson similarity.

## Data availability

All the metabolomics datasets described in our study are public datasets. MTBLS417 dataset was download from https://www.ebi.ac.uk/metabolights/MTBLS417 at 2019.12.29. 30STD_mix dataset was download from http://www.zhulab.cn/softwaredetail.php?id=40 at 2020.1.8.

## Code availability

All codes used in training, testing and evaluation are archived into DeepSWATH package, which is available at https://github.com/hcji/DeepSWATH under GPL (>= 3.0) license.

## Acknowledgements

This work is financially supported by the National Natural Science Foundation of China (Grant Numbers. 21873116 and 21675174)

## Additional information

### Supplementary information

Table S1. DeepSWATH processing results of 30 STD dataset.

Table S2. MS-DIAL processing results of 30 STD dataset.

Table S3. DDA MS/MS spectra obtained from MTBLS417 dataset by XCMS (positive mode).

Table S4. DDA MS/MS spectra obtained from MTBLS417 dataset by XCMS (negative mode).

Table S5. Precursor features of MTBLS417 dataset by XCMS (positive mode).

Table S6. Precursor features of MTBLS417 dataset by XCMS (negative mode).

Table S7. DeepSWATH processing results of MTBLS 417 dataset (positive mode).

Table S8. MS-DIAL processing results of MTBLS417 dataset (positive mode).

Table S9. DeepSWATH processing results of MTBLS417 dataset (negative mode).

Table S10. MS-DIAL processing results of MTBLS417 dataset (negative mode).

